# Bioprocess optimization enables enhanced protopanaxadiol production in yeast

**DOI:** 10.1101/2023.11.08.566192

**Authors:** Shangkun Qiu, Mariam Dianat Sabet Gilani, Conrad Müller, Ulf Liebal, Lars M. Blank

## Abstract

Ginsenosides are the primary active metabolites of ginseng and have been used in traditional herbal medicine in Asia for more than 4,000 years. Protopanaxadiol (PPD) is the precursor of dammarane-type ginsenosides, exhibiting different pharmacological activities. However, extraction of PPD from plant material is cumbersome because of its low concentration. Precision fermentation using recombinant yeast is a promising alternative strategy to produce PPD. For increasing PPD production, the medium and fermentation conditions were optimized by a Design of Experiment (DoE) approach. PPD production increased by 7.5-fold in the final cultivation condition compared to the reference condition. The PPD titer reached 1.2 g/L during simple 500 mL shake flask cultivations, the highest PPD production in shake flasks to date. The protocol presented facilitates parallel evaluation of recombinant yeast, thereby contributing to the much-needed sustainable synthesis of the versatile molecule class of triterpenoids.

## Introduction

Triterpenoids are secondary metabolites ubiquitous in plants with a range of biological properties.^1^ The triterpenoid class ginsenosides are mainly found in Ginseng (*Panax* spp.), a plant belonging to the Araliaceae family, which consists of 17 species, of which *Panax quinquefolius*, *Panax ginseng*, and *Panax notoginseng* are commonly industrially exploited. The plant extracts are discussed to possess pharmacological properties, including anti-cancer, anti-diabetic, anti-aging, anti-inflammatory, cardiovascular-protective, and the inhibition of coronavirus disease 2019 (COVID-19).^2,3,4^ More than 180 ginsenosides have been identified in the Panax genus, which can be categorized into two groups based on their aglycones: dammarane-type tetracyclic and oleanane-type pentacyclic ginsenosides.^5,6,7^ Dammarane-type tetracyclic ginsenosides can be further divided into three groups, ocotillol, protopanaxadiol (PPD)-type ginsenosides, and protopanaxatriol (PPT)-type ginsenosides, based on the carbohydrate moieties at C3, C6, and C20 positions and the respective hydroxylation.^7^

Ginseng has been used in traditional herbal medicine in Asia for more than 4,000 years.^5,8,3^ The term “ginseng” is derived from the Chinese words “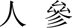 (Renshen)”, “the essence of men”, indicating its importance.^9^ Traditionally, ginsenosides were either consumed using the plant tissue or extracted from the root of ginseng. The root contains a documented total ginsenoside content of 60 mg/g in the dry weight.^3^ Changing and low amounts of the single target molecules in plant tissue make commercial production cumbersome.^10,11,8^ In addition, Ginseng roots need five to six years of growth until harvesting, with varying influences related to climate, land, soil, and pathogens.^3,5,12^ Furthermore, the complex structures of ginsenosides make chemical synthesis economically infeasible.^5,13^

Yeasts are simple eukaryotes that can be engineered for the synthesis of high-value metabolites. With their complete organelle structures, including mitochondria, peroxisomes, Golgi apparatus, endoplasmic reticulum (ER), and vacuoles, which allow post-translational modifications of proteins and operation of compartmented biochemical pathways, they are geared for enzymes from plants.^5,11,14,15^ Yeasts also possess redox systems that provide similar biological and physical environments encountered by cytochrome P450s and uridine diphosphate glycosyltransferase (UGTs) in plants and mammals.^5,14^ Due to these reasons, yeasts are considered excellent cell factories for producing natural plant products, such as ginsenosides, betulinic acid, farnesene, amorpha-4,11-diene, and others.^10,11,17,18,13^ For example, the production of α-farnesene was improved by engineered *Yarrowia lipolytica* during fed-batch fermentation via the expression of the acetyl-CoA acetyltransferase (Erg10p) from *Escherichia coli* and HMG-CoA reductase (Hmgp) from *Bordetella petrii*, the final α-farnesene production reached 26 g/L.^11,19^ Also, a highly engineered *Saccharomyces cerevisiae* expressing the amorpha-4,11-diene synthase (ADS) from *Artemisia annua* and using an unrestricted ethanol feed allowed the final production of 40 g/L amorpha-4,11-diene.^13^

Triterpenoids are either synthesized from squalene or 2,3-oxidosqualene, which are both intermediates of sterol synthesis in yeast.^5,16^ The synthesis of ginsenosides by engineered yeasts from renewable resources is a promising strategy to address the increasing demand for this interesting class of molecules.^5^ The microbial biosynthesis of ginsenosides has several advantages, including rapid growth, controllable culture conditions, space efficiency, sustainable production, environmentally friendly, and well-developed genetic manipulation technology.^5,11,20^ At present, only a fraction of ginsenosides are synthesized by engineered yeast, such as dammarenediol-II (DM), PPD, PPT, Compound K (CK), ginsenoside Rh1, ginsenoside Rh2, ginsenoside F1, ginsenoside Rg3, 20S-O-Glc-DM, 3β-O-Glc-DM, ginsenoside Rg1, ginsenoside NgR1, ginsenoside NgR2, ginsenoside Rf, ginsenoside Glc-Rf, and ginsenoside Ro.^5,21^ Furthermore, there have been some prominent achievements in synthesizing ginsenosides by yeasts. For example, according to a recent study, researchers constructed a PPD-producing *S. cerevisiae* cell factory via overexpression of mevalonate and PPD synthetic pathways and optimization of P450 expression levels; the final PPD production reached 11 g/L in 10 L fed-batch fermentation.^18^ Researchers also introduced the ginsenoside CK synthesis pathway into this strain while decreasing UDP-glucose consumption and improving UDP-glucose biosynthesis by engineering strategies; the final CK yield reached almost 6 g/L in fed-batch fermentations.^12^

Research showed that bioprocess optimization, including medium and cultivation conditions, can vastly improve the production of metabolites by cell factories.^22,23^ For example, carbon sources (e.g., glucose, glycerol, and glutamate), nitrogen sources, temperature, pH, oxygen, inoculum size, and salts significantly affect the cells’ growth and production.^22,24,25^ As the parameter space in medium and fermentation parameter optimization is huge, researchers aim for Design of Experiments (DoE) approaches. For example, researchers optimized the concentration of glucose, yeast extract, and peptone via the Box-Behnken design; this optimization increased the geranylgeraniol production by 1.75-fold compared to the traditional YPD medium.^26^ Cultured α-farnesene-producing *Y. lipolytica* in a medium optimized by a uniform design (UD) technique reached a final α-farnesene titer of 210 mg/L in a shake flask.^24^ In another study, adding citrate (0-10 mM) or acetate (0-10 mM) into the medium improved squalene production. The inhibition of the squalene monooxygenase Erg1p by terbinafine allowed 730 mg/L of squalene titers.^27^ Therefore, improving the cultivation conditions of engineered yeast is a promising strategy.

This study aims to provide a simple shake flask protocol to evaluate all the different mutants one might have to generate for further improving triterpenoid synthesis in yeast. Here, we use PPD synthesis as the main target, as the synthesis requires the activity of a triterpenoid cyclase and P450 monooxygenase. PPD is one of the 30-carbon dammarane II-type tetracyclic triterpenoids found in ginseng and is the precursor of high-value ginsenosides.^10^ PPD exhibits valuable biological properties, such as antineoplastic, antioxidant, antidepressant, antitumor, and anti-inflammation.^28,29^ For improved PPD production by yeast in shake flasks, the following parameters were considered: temperature, glucose and glutamate concentrations, salts solution, trace element solution, vitamin solution, and oxygen availability. PPD production reached 1.2 g/L using the best combination of shake flask set-up and optimized medium, the highest PPD production in shake flasks to date. This simple set-up facilitates the evaluation of engineered triterpenoid synthesizing yeast, and the amounts of triterpenoid produced enable advanced analytics and even first ideas for purification and use of this intriguing class of natural products.

## RESULTS AND DISCUSSION

### Effect of medium composition and Oxygen availability on DM and PPD Production

The DoE experiment included 34 runs designed using Design Expert 11 (Stat-Ease, Inc.). The seven significant factors and the high, medium, and low categories are shown in Table 1. The system Duetz microtiter plates were used for cultivation (Enzyscreen). Samples were taken after 72 h, 96 h, and 120 h of cultivation. DM is the precursor of PPD, which is also a triterpenoid and is the substrate of the cytochrome P450 monooxygenase protopanaxadiol synthase (PPDS) for PPD (Figure 1).^30,31^ DM and PPD concentrations were highest after 120 h. PPD production reached 0.3 g/L in four experiments (1, 10, 12, and 33) (Figure 2A). Among them, runs 1, run 10, and run 12 were all cultured at 20 ℃, and run 33 was cultured at 30 ℃. Glucose and glutamate concentrations were three-fold compared to standard WM8+ medium (100 g/L) for runs 1, 12, and 33. Notably, the salt concentration of all four runs was 0.33-fold of standard WM8+ medium. Also, higher oxygen availability was common in these four experiments, using only 1 mL of culture broth. The PPD concentration reached 0.9 g/L in run 1, the highest PPD production among all 34 runs in the DoE experiment.

**Figure 1.**
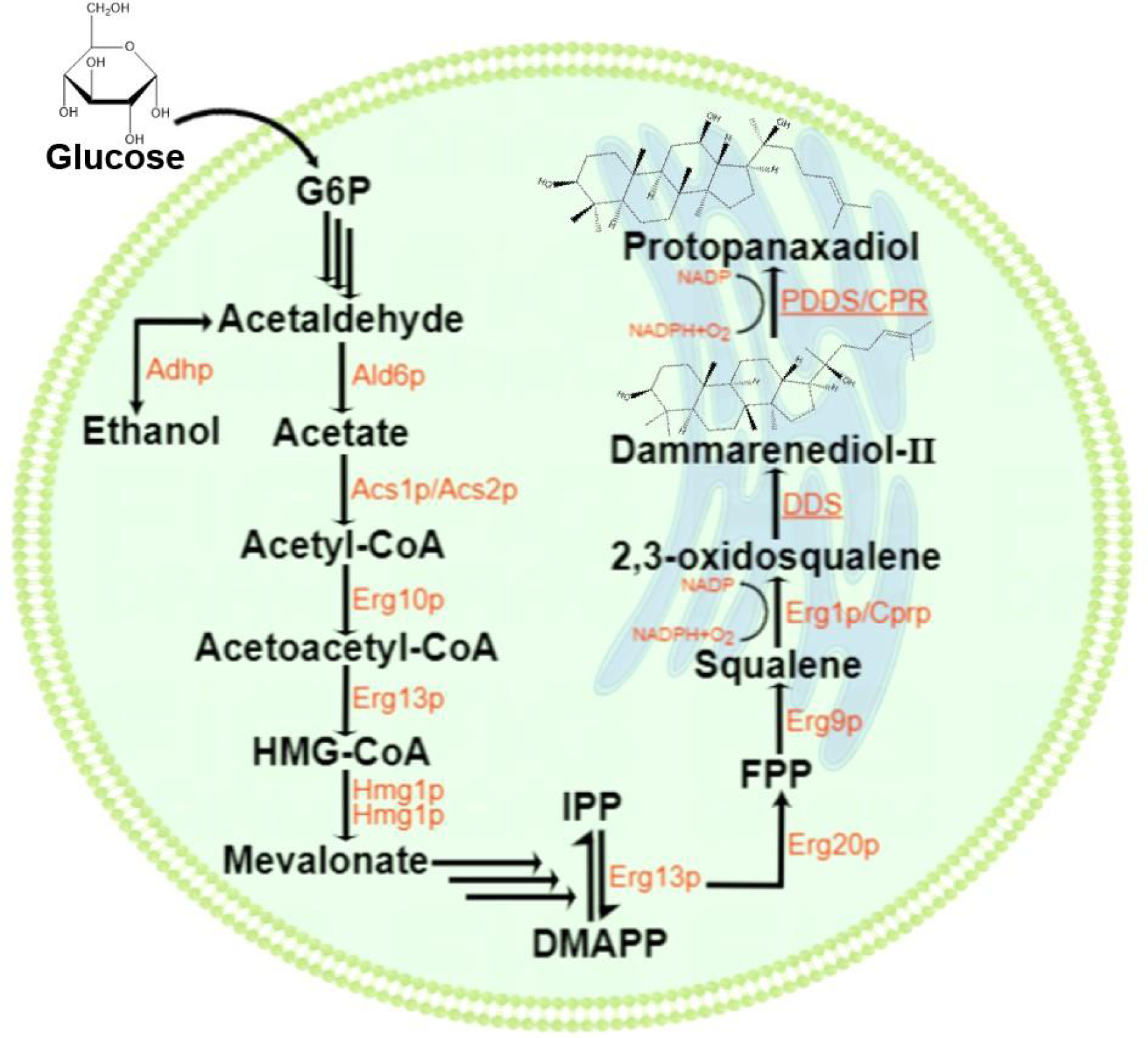
The PPD synthesis pathway in engineered *S. cerevisiae*. The recombinant enzymes are underlined. G6P, glucose 6-phosphate; HMG-CoA, 3-hydroxy-3-methylglutaryl-CoA; IPP, isopentenyl pyrophosphate; DMAPP, dimethylallyl diphosphate; FPP, farnesyl diphosphate; Erg10p, acetyl-CoA acetyltransferase; Erg13p, HMG-CoA synthase; Hmg1p and Hmg2p, HMG-CoA reductase 1 and 2; Idi1p, isopentenyl-diphosphate delta-isomerase; Erg20p, farnesyl diphosphate synthase; Erg9p, squalene synthase; Erg1p, squalene monooxygenase; DDS, dammarenediol-II synthase; PPDS, protopanaxadiol synthase; ER, endoplasmic reticulum. This image was created by Figdraw.

**Figure 2.**
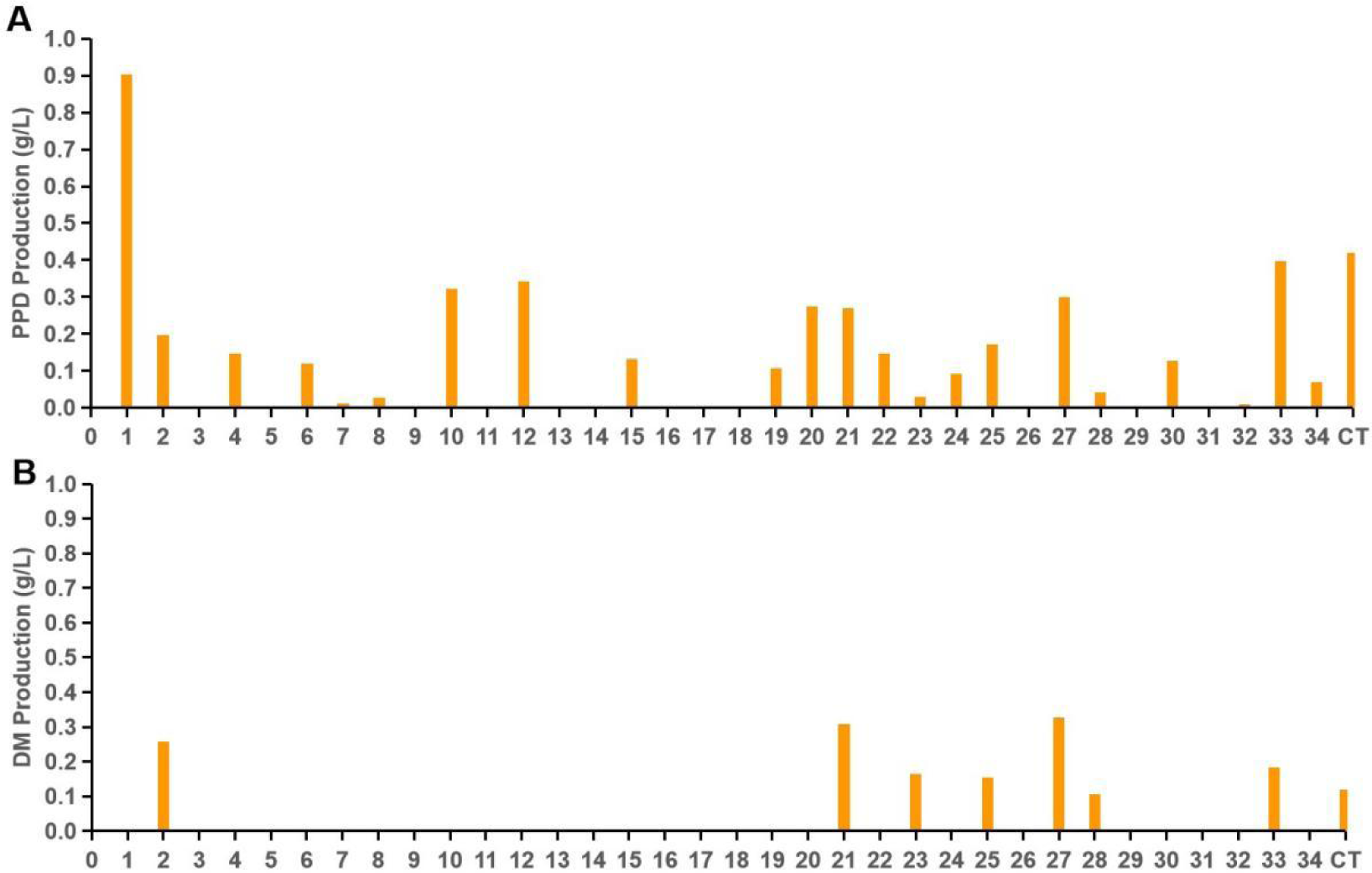
The result of the 34 runs DoE experiment. (A) PPD production by engineered yeast in 24-well plates. (B) DM production at different fermentation conditions in 24-well plates. CT: Control experiment, standard WM8+ medium.

**Table 1.** DoE for improving the standard WM8+ medium and fermentation conditions.

| Run | Factor A:<br>Glucose | Factor B:<br>Salts | Factor C:<br>Glutamate | Factor D: Trace<br>element | Factor E:<br>Vitamin | Factor F:<br>Temperature [°C] | Factor G:<br>Volume [mL] |
| --- | --- | --- | --- | --- | --- | --- | --- |
| 1 | 3 | 0.33 | 3 | 3 | 0.33 | 20 | 1 |
| 2 | 1.67 | 1.67 | 1.67 | 1.67 | 1.67 | 25 | 2 |
| 3 | 3 | 3 | 0.33 | 3 | 0.33 | 30 | 3 |
| 4 | 3 | 3 | 0.33 | 0.33 | 3 | 30 | 3 |
| 5 | 0.33 | 0.33 | 0.33 | 3 | 0.33 | 20 | 3 |
| 6 | 3 | 3 | 3 | 3 | 3 | 20 | 1 |
| 7 | 3 | 3 | 0.33 | 3 | 0.33 | 20 | 1 |
| 8 | 0.33 | 0.33 | 3 | 0.33 | 0.33 | 30 | 1 |
| 9 | 0.33 | 0.33 | 0.33 | 0.33 | 3 | 30 | 1 |
| 10 | 0.33 | 0.33 | 3 | 3 | 3 | 20 | 1 |
| 11 | 0.33 | 3 | 3 | 0.33 | 3 | 30 | 1 |
| 12 | 3 | 0.33 | 3 | 0.33 | 3 | 20 | 1 |
| 13 | 0.33 | 3 | 3 | 3 | 0.33 | 20 | 3 |
| 14 | 0.33 | 3 | 3 | 0.33 | 3 | 20 | 3 |
| 15 | 0.33 | 0.33 | 0.33 | 3 | 0.33 | 30 | 1 |
| 16 | 0.33 | 3 | 0.33 | 0.33 | 0.33 | 30 | 3 |
| 17 | 3 | 3 | 0.33 | 0.33 | 0.33 | 30 | 1 |
| 18 | 0.33 | 0.33 | 3 | 0.33 | 0.33 | 20 | 3 |
| 19 | 0.33 | 3 | 0.33 | 3 | 3 | 30 | 1 |
| 20 | 3 | 0.33 | 3 | 3 | 3 | 20 | 3 |
| 21 | 3 | 3 | 3 | 0.33 | 0.33 | 30 | 3 |
| 22 | 3 | 0.33 | 0.33 | 3 | 3 | 20 | 1 |
| 23 | 1.67 | 1.67 | 1.67 | 1.67 | 1.67 | 25 | 2 |
| 24 | 0.33 | 3 | 0.33 | 0.33 | 3 | 20 | 1 |
| 25 | 1.67 | 1.67 | 1.67 | 1.67 | 1.67 | 25 | 2 |
| 26 | 3 | 0.33 | 0.33 | 0.33 | 0.33 | 30 | 3 |
| 27 | 3 | 3 | 3 | 3 | 3 | 30 | 3 |
| 28 | 3 | 3 | 0.33 | 0.33 | 0.33 | 20 | 3 |
| 29 | 0.33 | 0.33 | 3 | 3 | 0.33 | 30 | 3 |
| 30 | 3 | 0.33 | 0.33 | 0.33 | 3 | 20 | 3 |
| 31 | 0.33 | 0.33 | 0.33 | 3 | 3 | 30 | 3 |
| 32 | 0.33 | 3 | 3 | 0.33 | 0.33 | 20 | 1 |
| 33 | 3 | 0.33 | 3 | 3 | 3 | 30 | 1 |
| 34 | 2 | 1.67 | 1.67 | 1.67 | 1.67 | 25 | 2 |
| CT | 1 | 1 | 1 | 1 | 1 | 30 | 2 |

Every factor has a low (red), middle (yellow), and high (green) value. For factors A-E, the figures in the table represent fold changes in the actual concentration compared to the standard WM8+ medium. For factor F, the figures in the table represent the actual temperature. For factor G, figures in the table represent the volume of the medium in 24-well plates. CT: Control experiment, standard WM8+ medium.

DM production suggests triterpenoid synthesis. However, it is also a potential limitation in the subsequent P450-catalyzed reaction to PPD. Only seven experiments showed DM production (Figure 2B). Among the seven experiments, only runs 21 and 27 had DM concentrations above 0.3 g/L. These two runs’ glucose, glutamate, and salt concentrations were three-fold that of standard WM8+ medium. The results suggest high glucose and glutamate concentrations support high DM and PPD production. At the same time, low salt concentration, low temperature, and high oxygen concentration might benefit PPD production, indicating that the P450-catalyzed hydroxylation is somewhat challenging, with substrate limitation and a slow rate of catalysis.

### The ordinary least squares (OLS) Estimation of the DoE Data

DoE data was analyzed with OLS regression to PPD and DM to identify the predictive factors. We tested models containing only the individual factor effects (Ind-OLS) as well as individual factors combined with two-factor interactions (Int-OLS) (Table 2). The p-value of the F-Statistics identified models with sufficient power to explain the variance of the observations (PPD) if the p-value is below 0.05. The Ind-OLS and the more complex Int-OLS explain 41% and 83% of the variance in PPD, respectively (adj. R²(Ind-OLS)=0.41, adj. R²(Int-OLS)=0.83). The p-values of the individual factors in each regression model determine whether the factor significantly contributes to the prediction. There are two significant factors for Ind-OLS (A: glucose, C: glutamate) and seven for Int-OLS (B: salt, C: glutamate, AxE (vitamins), AxC, B (salt)xF (temperature), CxD (trace element), BxG (oxygen)) (Figure 3). In the simple Ind-OLS model with individual factors, only glucose and glutamate are the most influential predictors that positively contribute to PPD production with correlation values of 0.4 (Figure 3A). Conversely, salt concentration (B), temperature (F), and medium volume (G) correlate negatively to the final PPD production, albeit with low correlation values at −0.2 (Figure 3A). In the more complex Int-OLS model, glucose is only significant via the two-factor interactions with glutamate (A:C) and vitamin (A:E) but no longer as an individual factor. The most significant single factors in the complex Int-OLS model are the positive correlation with glutamate (C) and the negative correlation with salts (B). While the volume level (G, oxygen) individually is just above the significance value (p=0.05), it negatively correlates with PPD and is also negative with the significant interaction with salts (BxG). Thus, less oxygen via higher volumes decreases PPD (Figure 3A). The yeast increases biomass quickly, while the oxygen concentration decreases fast in the exponential phase.^25^ Furthermore, NADPH is oxidized to NADP by a P450 reductase in the presence of oxygen, then transfers electrons to the monooxygenases, such as PPDS.^7^ Therefore, controlling the oxygen concentration will also play a critical role in synthesizing the final product. Consequently, in the subsequent investigation, we selected glucose, glutamate, salts, temperature, and oxygen as the main significant factors for further research to check if these factors will affect the final PPD and DM production.

**Figure 3.**
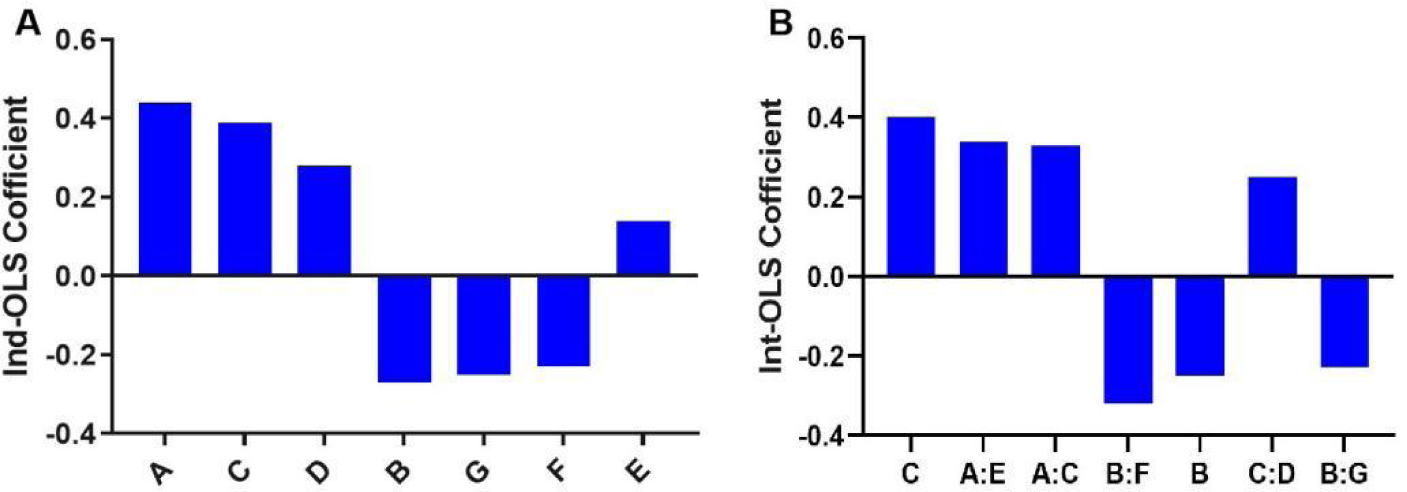
The OLS regression result of the DoE data with prediction of PPD production. (A) The OLS regression result of the Ind-OLS. (B) The OLS regression result of the Int-OLS. The coefficients with decreasing absolute values are shown for factors (Ind-OLS) and factor interactions (Int-OLS), significantly explaining the variance (p-values below 0.05). The larger the coefficient, the greater the influence of this factor or factor interaction on the final PPD production. A: glucose; B: salts; C: glutamate; D: trace element solution; E: vitamin solution; F: temperature; G: medium volume.

**Table 2.** Regression results for OLS with seven factors alone and first-order interactions of two parameters.

| Observable | Equation | F-Statistic<br>p-value | Adj. R<sup>2</sup> |
| --- | --- | --- | --- |
| PPD | Ind-OLS: $\sum c_F F$ | 0.003 | 0.41 |
| DM |  | 0.14 | 0.14 |
| PPD | Int-OLS: $\sum c_F F + \sum c_{Fij} F_i F_j$ | 0.02 | 0.83 |
| DM |  | 0.83 | -0.53 |

The F-Statistic shows the p-value for the hypothesis that the independent variables of the OLS model explain the variability of the dependent variable, R<sup>2</sup> is the percentage of explained variance, Adj. R<sup>2</sup> is a multi-variable corrected correlation coefficient, Fact. with p<0.05, are factors with a p-value below 0.05, along with the estimated coefficient in the OLS model.

### Effect of Temperature on DM and PPD Production

Sesquiterpenoid production by engineered *S. cerevisiae* was reported to be better at 25 °C than at 30 °C.^32^ We also investigated temperature as an optimization parameter for PPD production. Here, the highest PPD production was observed at 20 ℃ in the 34 runs DoE experiment (Figure 2A). Temperatures above 30 °C were reported to cause stress to bakeŕs yeast and decrease cell viability.^25^ Therefore, the 15 runs of the DoE experiment cultured at 20 ℃ were selected to investigate the effect of different temperatures on DM and PPD production in 24-well plates. To explore the optimum temperature, 18 ℃, 20 ℃, 22 ℃, 25 ℃, 28 ℃, 30 ℃, and 32 ℃ were selected. The other fermentation and medium conditions remained constant. In general, the production of PPD increased with increasing temperature and reached the highest value at 28°C, then decreased with increasing temperature (Figure 4A, Figure S1). Especially the PPD productions of runs 1 and 12 are the highest among the 15 runs, reached 1.5 g/L at 28 ℃ (Figure 4A). But for run 13, run 14, run 18, run 20, run 24, run 30, and run 32, the PPD production reached the highest value at 32 ℃. However, the PPD production for these seven runs is lower than runs 1 and 12 (Figure S1). So, we can speculate that the optimum temperature for PPD production is 28 ℃.

**Figure 4.**
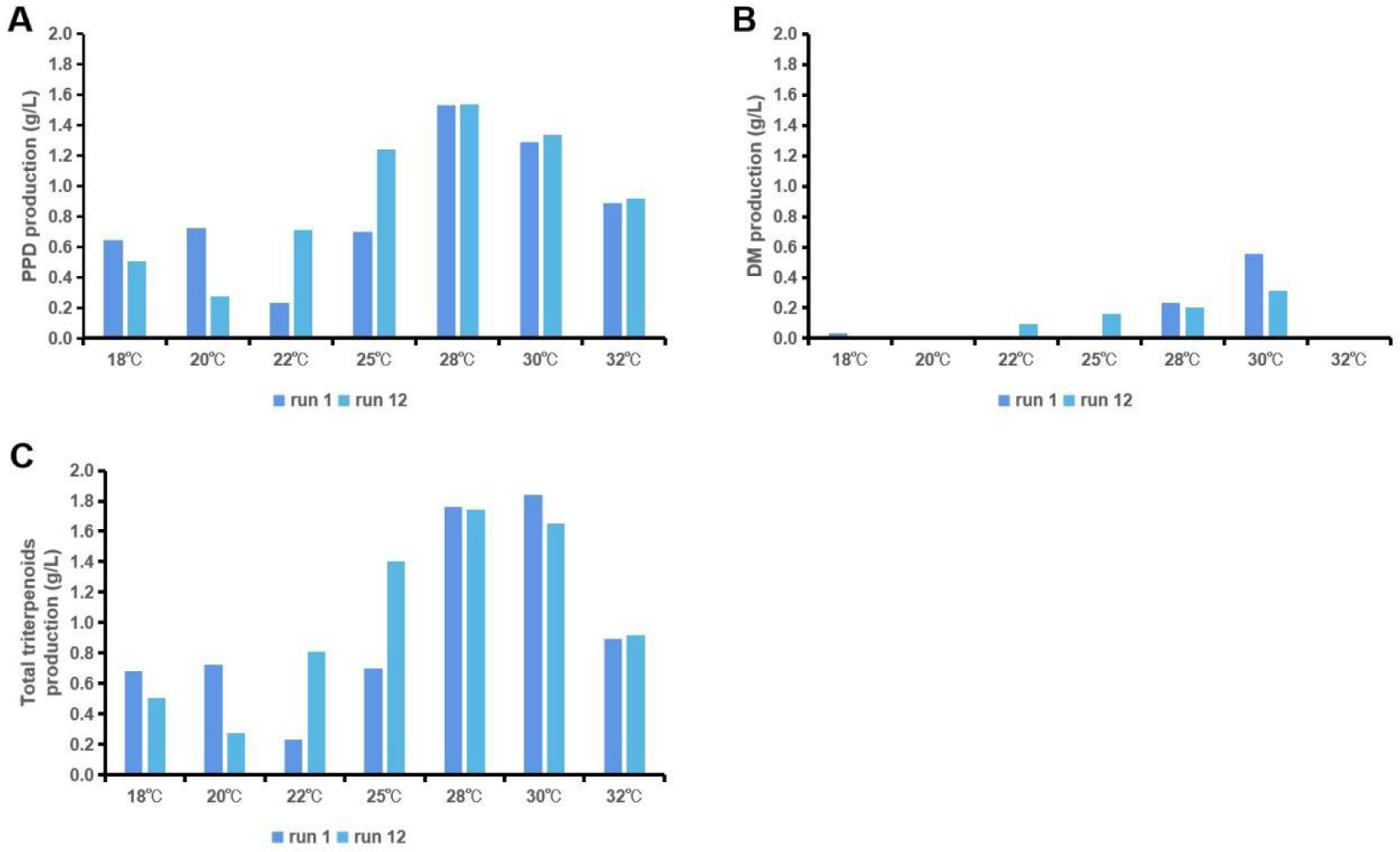
The effect of different temperatures on DM and PPD production. (A) The comparison of PPD production between run 1 and run 12 at 18℃, 20℃, 22℃, 25℃, 28℃, 30℃, and 32℃. (B) The comparison of DM production between run 1 and run 12 at 18℃, 20℃, 22℃, 25℃, 28℃, 30℃, and 32℃. (C) The comparison of total triterpenoid production between run 1 and run 12 at 18℃, 20℃, 22℃, 25℃, 28℃, 30℃, and 32℃. Other runs’ data is shown in Figure S1.

The DM production of run 1, run 12, and run 24 reached the highest when cultured at 30 ℃. However, the optimum temperature of the run 6 and run 32 is 25 ℃, run 10, run 20, run 22, and run 28 is 28 ℃, and run 30 is 32℃ (Figure 4B, Figure S1). The first reason why the optimum temperatures of PPD and DM synthesis in all the runs are different may be that the different concentrations of compounds in the medium can affect the DM conversion to PPD at different temperatures. Another reason may be that the different medium compositions affect the enzyme activities involved in PPD and DM synthesis at different temperatures. The DM production of run 28 is the highest when cultured at 28 ℃, which reached 1.1 g/L (Figure S1). The DM production of run 22, run 28, and run 30 are all over 0.8 g/L. The total triterpenoid production of run 1 is the highest among all the runs, cultured at 30 ℃ and reached 2.3 g/L (Figure 4C). The total triterpenoid production of run 1, run 12, and run 30 is over 2 g/L. However, the total triterpenoid production of run 1 and run 12 reached 2 g/L when cultured at 28 ℃ or 30 ℃, while run 30 reached 2 g/L at 32 ℃ (Figure 4C, Figure S1). Hence, optimum temperature for DM and total triterpenoid production is in the range of 28-32 ℃.

### Effect of Glucose, Glutamate, and Salt Concentration on DM and PPD Production

According to the results above, runs 1 and 12 have the highest PPD and total triterpenoid production. The conditions of runs 1 and 12 were chosen for further investigation (Figure 4A). Glucose, glutamate, and salt were selected to be varied in concentration, while all other parameters were kept constant. The optimum salt concentration for PPD production seems to be 0.33, 0.44 or 2 times the WM8+ medium, with the highest PPD production is 2 g/L (Figure 5A). Here, it is indicated that the optimum concentration of glucose and glutamate for PPD synthesis is two times that of the standard WM8+ medium. Higher concentrations might cause osmotic stress and deflect resources from PPD production (Figure 5A).

**Figure 5.**
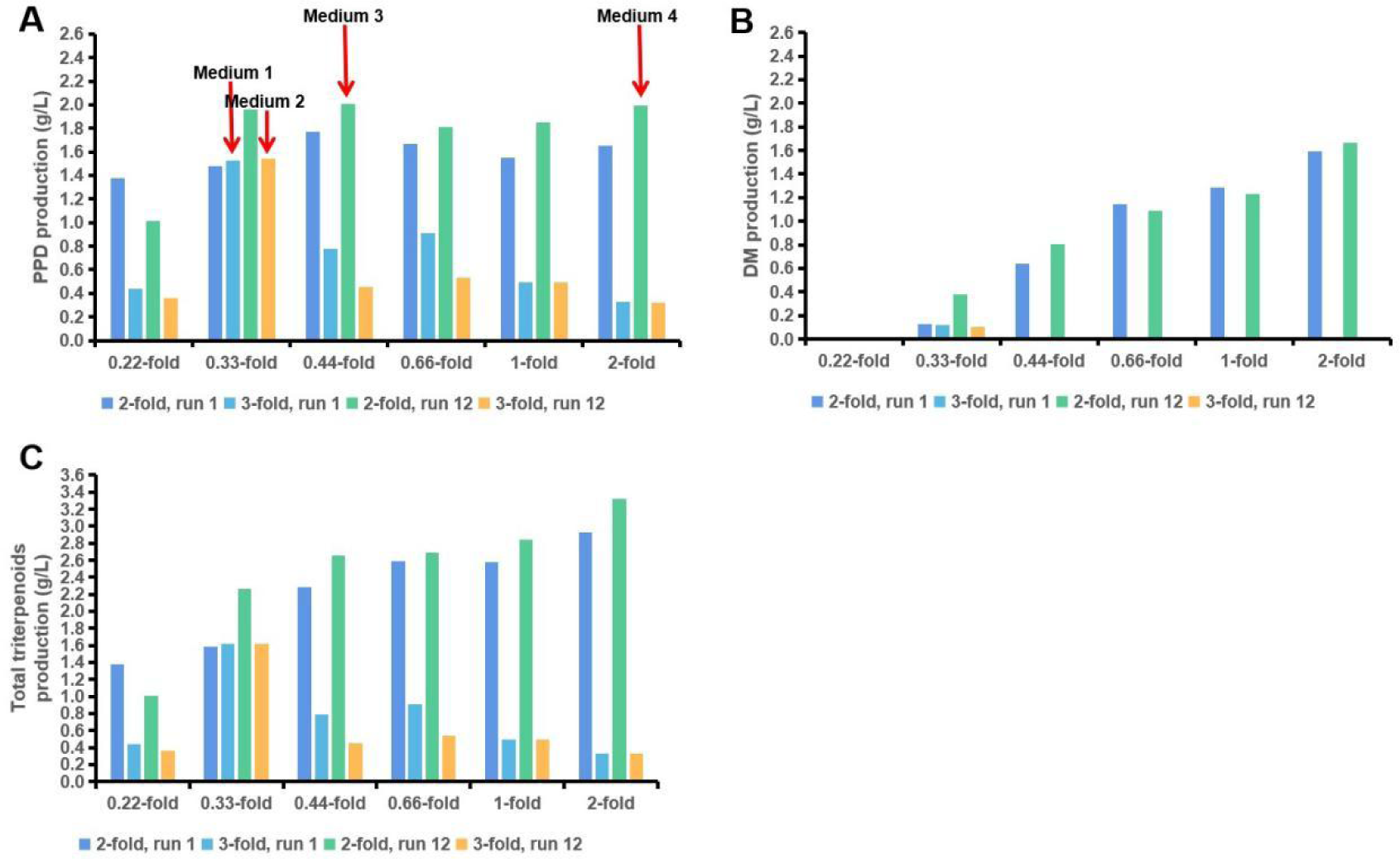
The effect of glucose, glutamate, and salt concentration on PPD and DM production (A) PPD production depends on glucose, glutamate, and salts. (B) DM production is dependent on glucose, glutamate, and salts. (C) The total triterpenoid production. The glucose and glutamate concentration is two or three times the standard WM8+ medium. The horizontal coordinate is the salt concentration compared to the standard WM8+ medium. Runs 1 or 12 means the medium is modified from run 1 or run 12.

DM production profits from increased salt concentration, and this is different from PPD. Again, three times or even higher concentrations of glucose and glutamate abolish DM production. The highest DM production of all the runs was 1.6 g/L (Figure 5B). Indeed, under these conditions, doubling the glucose and glutamate concentrations of WM8+ medium, total triterpenoid production exceeded 3 g/L (Figure 5C).

For the translation of intracellular products, the yield per biomass is central, as is the yield on substrate for any biotechnological process. Here, biomass generally increased with increasing salt concentrations under constant carbon sources (Figure 6). The highest biomass yield per gram carbon source is 328 mgCDW/g_glucose and glutamate_ in two-fold salt concentration (Figure 6C). The triterpenoid yield increased with increasing salt concentrations when glucose and glutamate concentration were twice that of the standard WM8+ medium. However, the PPD yield first increases and then decreases with increasing salt concentrations (Figure 6A and C). The highest total triterpenoid yield is 38 mg/g_glucose and glutamate_ (Figure 6C). At the same time, the highest PPD yield is 21 mg/g_glucose and glutamate_ (Figure 6C).

**Figure 6.**
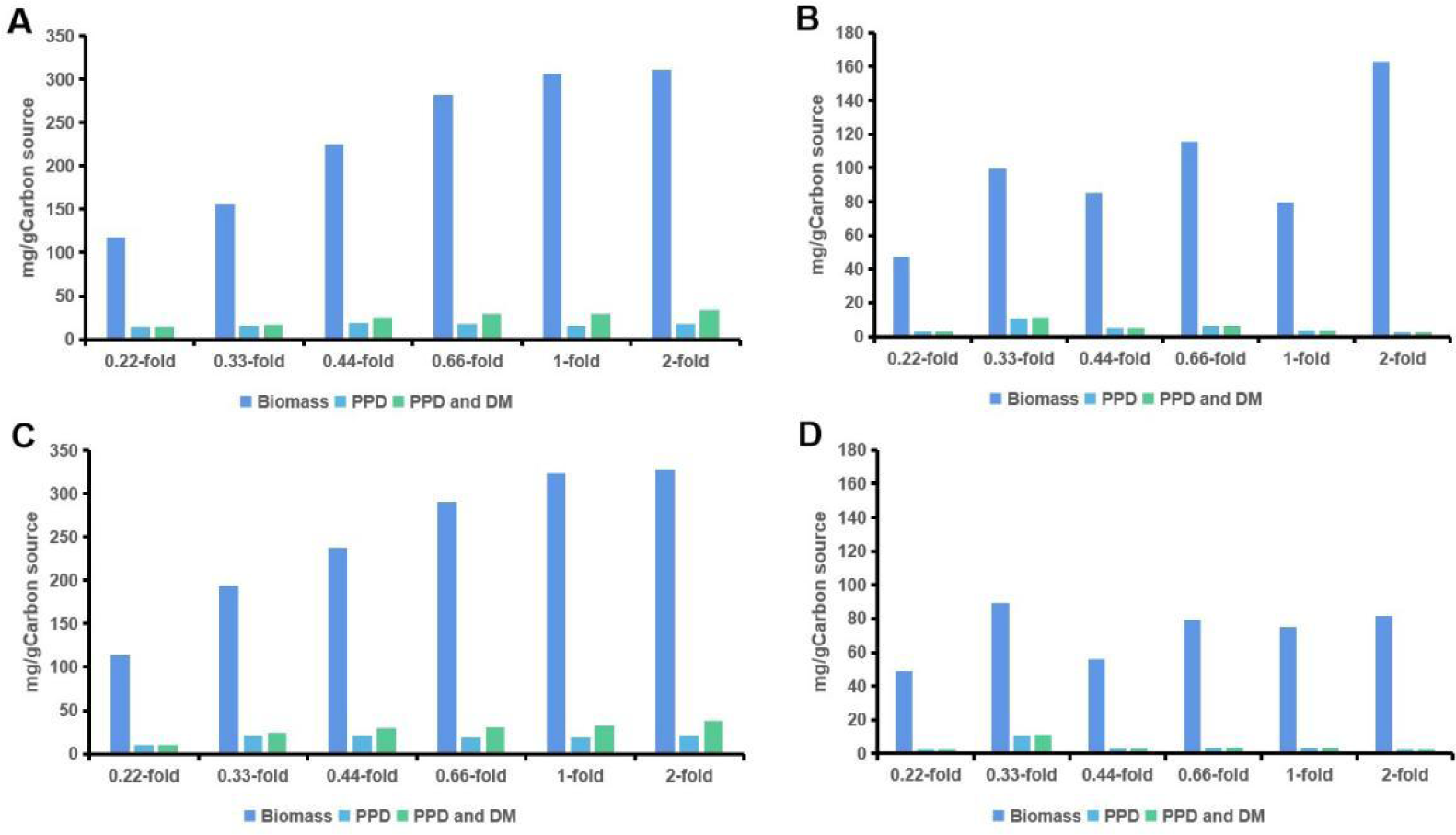
The effect of glucose, glutamate, and salt concentration on triterpenoid yield. (A) The glucose and glutamate concentration is two times the standard WM8+ medium. This medium is modified from run 1. (B) The glucose and glutamate concentration is three times the standard WM8+ medium. This medium is modified from run 1. (C) The glucose and glutamate concentration is twice the standard WM8+ medium. This medium is modified from run 12. (D) The glucose and glutamate concentration is three times the standard WM8+ medium. This medium is modified from run 12. The horizontal coordinate is the salt concentration compared to the standard WM8+ medium. All other parameters were used as described in Figure 5.

The total triterpenoid content of ripe ginseng root is 0.1-3 %.^3^ We here estimated the CDW content of PPD, DM, and total triterpenoids. The CDW content of the total triterpenoid of three runs is above 120 mg/g_CDW_ (Figure 7A and C). The carbon source concentration of these three runs is two times that of the standard WM8+ medium. The salt concentration is 0.22, 0.33, or 0.44 times the reference. All the data indicates that low salt concentrations favor high CDW content of PPD. In contrast, high salt concentrations can contribute to increased CDW content of DM (Figure 7A and C). The optimum glucose and glutamate concentration for the CDW content of PPD and total triterpenoids is two times of the standard WM8+ medium.

**Figure 7.**
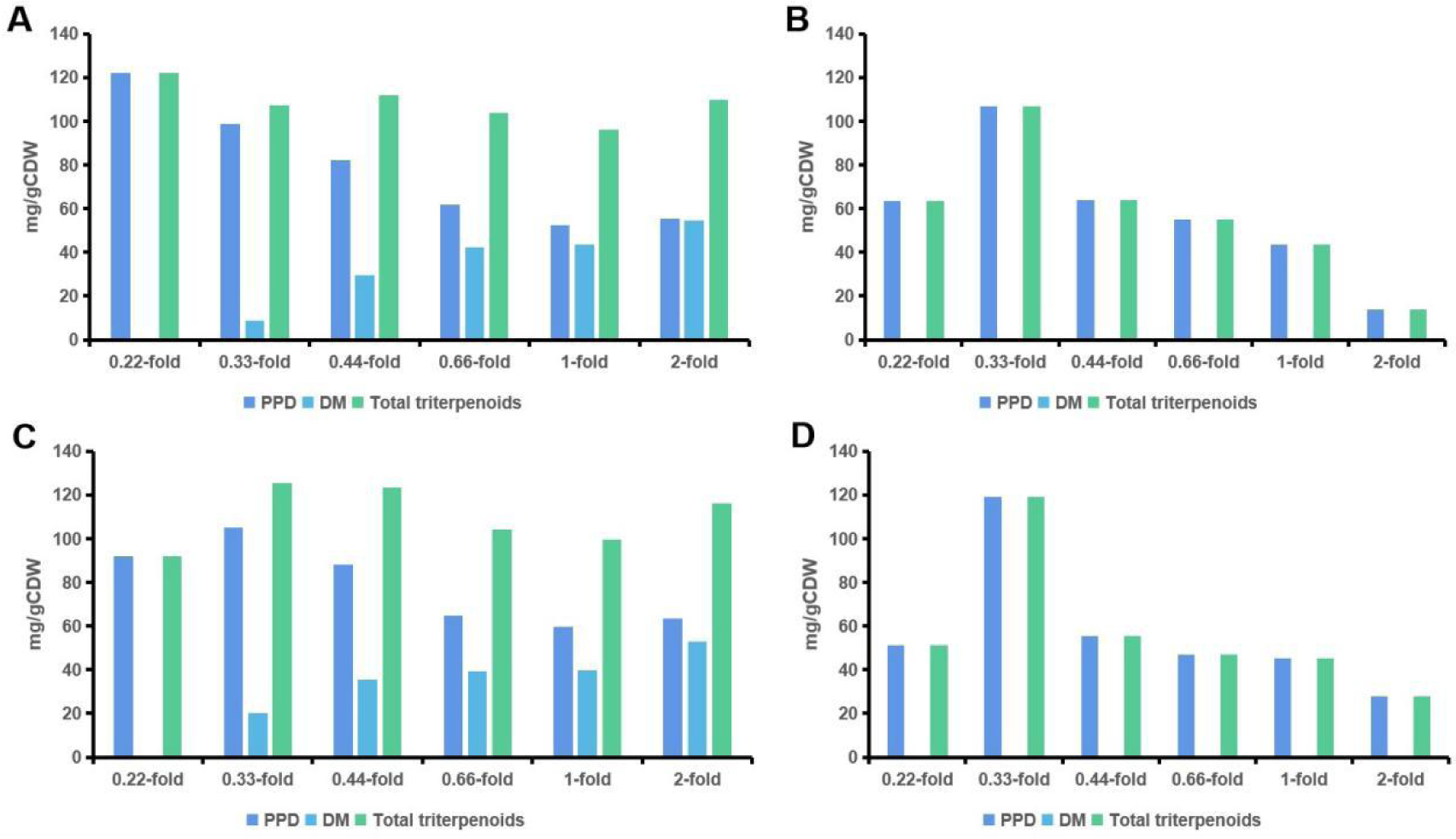
The effect of glucose, glutamate, and salt concentration on CDW content of triterpenoids. (A) The glucose and glutamate concentration is two times the standard WM8+ medium. This medium is modified from run 1. (B) The glucose and glutamate concentration is three times the standard WM8+ medium. This medium is modified from run 1. (C) The glucose and glutamate concentration is twice the standard WM8+ medium. This medium is modified from run 12. (D) The glucose and glutamate concentration is three times the standard WM8+medium. This medium is modified from run 12. The horizontal coordinate is the salt concentration compared to the standard WM8+ medium. All other parameters were used as described in Figure 5.

### Triterpenoid Production in 500 mL Shake Flasks

While microtiter plate cultivation allows strain comparison, shake flask cultivations can be designed to test product purification ideas and synthesize the first products for use. For this scale increase, we selected four media from the above experiments, which are medium 1 (3-fold, run 1), medium 2 (3-fold, run 12), medium 3 (2-fold, run 12), and medium 4 (2-fold, run 12) (Figure 5A). Yeasts were cultivated in two types of 500 mL shake flasks. The shake flask was closed by either an aluminum or a membrane lid (Figure 8A). The membrane lid facilitates better aeration when compared to the aluminum lid. Two enzymes in PPD synthesis require oxygen: the conversion of squalene to 2,3-oxidosqualene, catalyzed by Erg1p, and the conversion of DM to PPD, catalyzed by PPDS (Figure 1).^33,34^ First, yeasts were cultivated in the shake flask with an aluminum lid. The smaller the filling volume of the shake flask, the better the aeration. Two different filling volumes were used, i.e., 25 mL and 50 mL. The yeast was cultured in the four improved media. The results showed the highest PPD production of 0.8 g/L occurred in medium 3 with 25 mL filling volume. The PPD production decreased with the increase in the medium. The PPD production in all four media tested is higher than that in WM8+ medium, which is 0.16 g/L. It can be speculated that the higher oxygen concentration benefits the synthesis of PPD, which can increase the final PPD production in the shake flask (Figure 8B).

**Figure 8.**
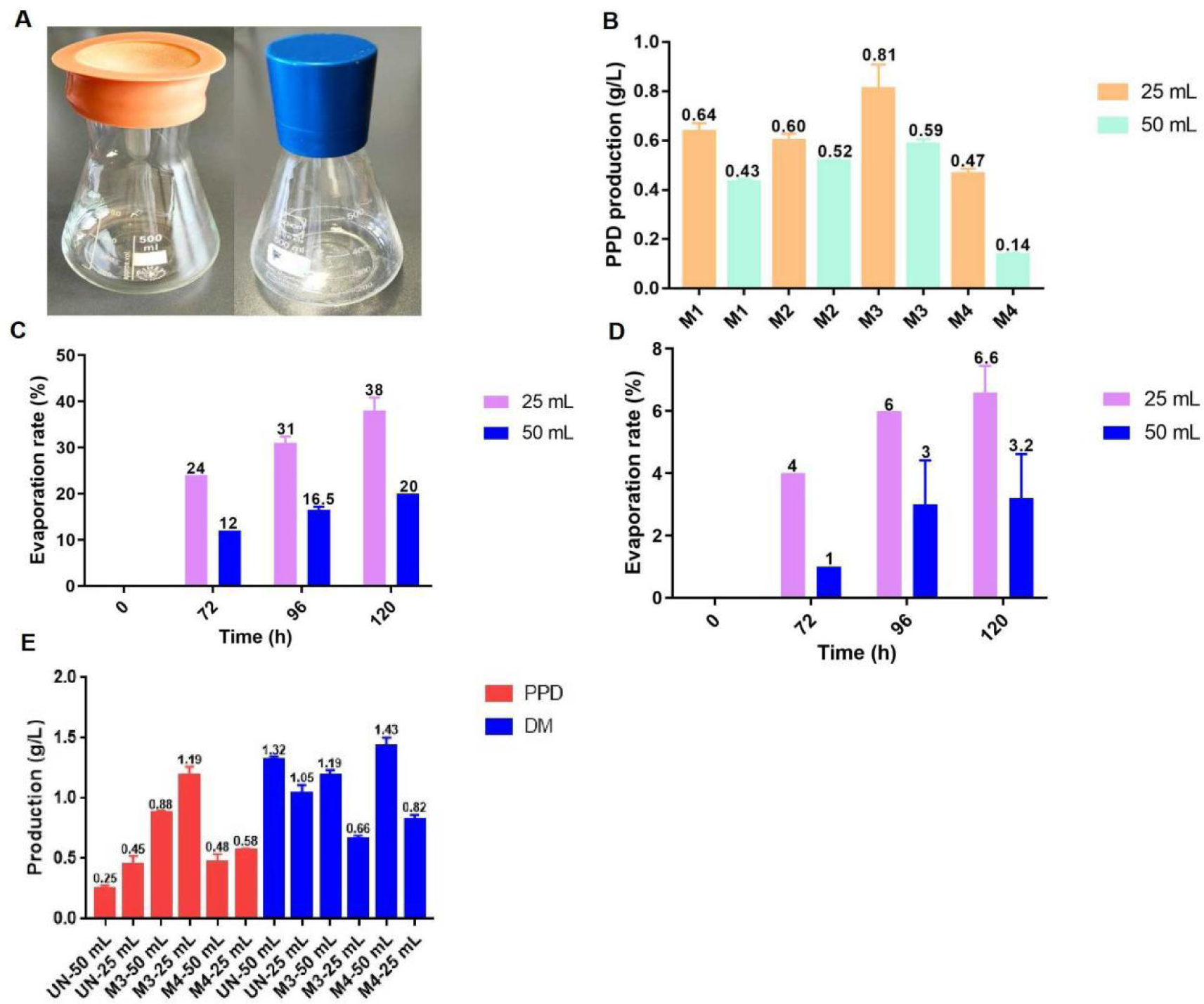
Shake flask configuration influences PPD production. (A) Shake flasks with membrane and aluminum lid. (B) PPD production depends on the filling volume using a shake flask with an aluminum lid. (C) Water evaporation rate in the 500 mL shake flask with a membrane lid. (D) Water evaporation rate in the 500 mL shake flask with an aluminum lid. (E) PPD and DM production depends on the filling volume using a shake flask with a membrane lid. UN, unoptimized standard WM8+ medium; M1, Optimum medium 1; M2, Optimum medium 2; M3, Optimum medium 3; M4, Optimum medium 4.

The identical experiments were repeated using the membrane lid for improved aeration. Notably, water evaporation is higher using the membrane lid (Figure 8C and D), although the shaker was equipped with a moisture controller, keeping water content in the air at 70% saturation. To account for evaporation, we calculated the water evaporation in the 500 mL shake flask with both types of lids at 28 ℃ (Figure 8C and D). We measured the volume after 120 h and calculated the product concentration without evaporation. The corrections were as follows:

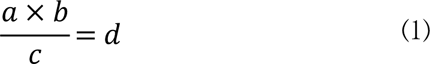

where *a* refers to the PPD or DM production (g/L) measured by HPLC; *b* refers to the final volume (mL); *c* refers to initial volume (mL); *d* refers to the actual PPD or DM production (g/L) without evaporation.

The results indicated that the PPD production also decreased with the increase of medium volume. The PPD productions of optimum media 3 and 4 in this membrane lid shake flask are higher than the PPD production of the same media with an aluminum lid, reaching 1.2 g/L and 0.6 g/L when the volume is 25 mL. At the same time, the DM production increased with medium volume (Figure 8E). This indicated that the membrane lid and smaller volume allow more oxygen to enter the shake flask, which supplies more oxygen for improving the conversion of squalene to 2,3-oxidosqualene and the conversion of DM to PPD (Figure 1). The highest PPD productions of unoptimized standard WM8+ medium in the shake flask with an aluminum lid or membrane lid are 0.16 g/L or 0.45 g/L, respectively (Figure 8E). This indicated that the highest PPD production of the optimum medium 3 is 2.6-fold compared to the unoptimized standard WM8+ medium in the shake flask with a membrane lid. However, the PPD productions in media 1 and 2 with this membrane lid shake flask are much lower than the PPD production in medium 3 (data not shown). This may be due to the higher glucose and glutamate concentration (three times the standard WM8+ medium) in media 1 and 2, limiting the PPD and DM synthesis in the shake flask, just like the analysis described in Figure 5.

### The fermentation kinetics of PPD and DM in engineered yeast cultured in the standard WM8+ medium and the M3 medium

Ethanol is a more reduced carbon source than glucose, which feeds directly into the cytosolic acetyl-CoA pool required for the mevalonate pathway.^7^ Therefore, we also measured the concentration of the carbon sources and triterpenoid products during the fermentation to investigate the fermentation kinetics of PPD and DM. As a Crabtree-positive yeast, the engineered *S. cerevisiae* uses respiro-fermentative glucose metabolism and produces ethanol and carbon dioxide.^35^ After glucose depletion at 24 h, ethanol will be the carbon source in the presence of oxygen, as the classic diauxic shift of *S. cerevisiae*. The ethanol concentration also reached the highest value at 24 h, 12.5 g/L, and 35 g/L in the standard WM8+ and M3 medium (Figure 9A and B). Ethanol has been shown to be a beneficial carbon source for triterpenoid production.^7^ Therefore, PPD and DM synthesis was very fast when the carbon source became ethanol. The highest PPD production is 0.6 g/L and 1.8 g/L in the standard WM8+ and M3 medium, respectively, slightly higher than the PPD production above data (0.45 g/L in the standard WM8+ medium and 1.2 g/L in the M3 medium) (Figure 8E, 9A and B). The reason should be due to the several samples taken during the fermentation that reduced the volume and increased the oxygen concentration; this led to higher PPD production. The result indicated that the M3 medium performs better than the standard WM8+ medium for PPD synthesis.

**Figure 9.**
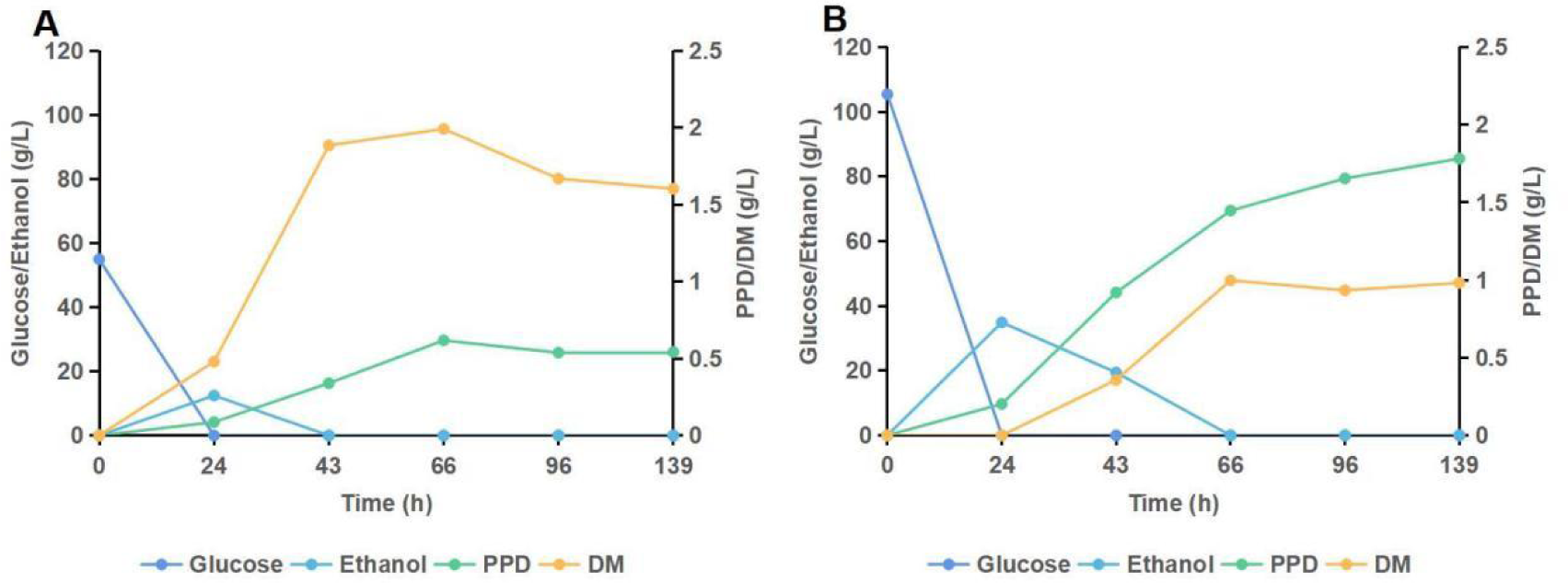
Fermentation kinetics of PPD and DM in engineered yeast. (A) The concentration of glucose, ethanol, PPD, and DM during the fermentation in engineered yeast cultured in the standard WM8+ medium. (B) The concentration of glucose, ethanol, PPD, and DM during the fermentation in engineered yeast cultured in the M3 medium.

According to previous studies, it is a promising choice to improve the production of the final metabolites in the cell’s factories by optimizing the fermentation bioprocess, including medium compounds and fermentation conditions.^22,23^ The PPD production of the M3 medium is nearly 2 g/L in the 24-well plates, which is much more than the PPD production in the shake flask with a membrane lid, 1.2 g/L. The reasons may be due to the cells’ sediment, the water evaporation, some cells on the wall, oxygen concentration, and other factors that are different between the two growth formats. We also checked other engineered yeasts that can produce other triterpenoids (betulin and betulinic acid) with the M3 medium. The result showed that the betulin and betulinic acid productions of these two strains in the M3 medium are largely increased compared to the productions in the standard WM8+ medium (Figure 10). This means that the M3 medium also works for other engineered strains. As aimed, the results can be generalized to evaluate engineered yeasts under simple growth conditions.

**Figure 10.**
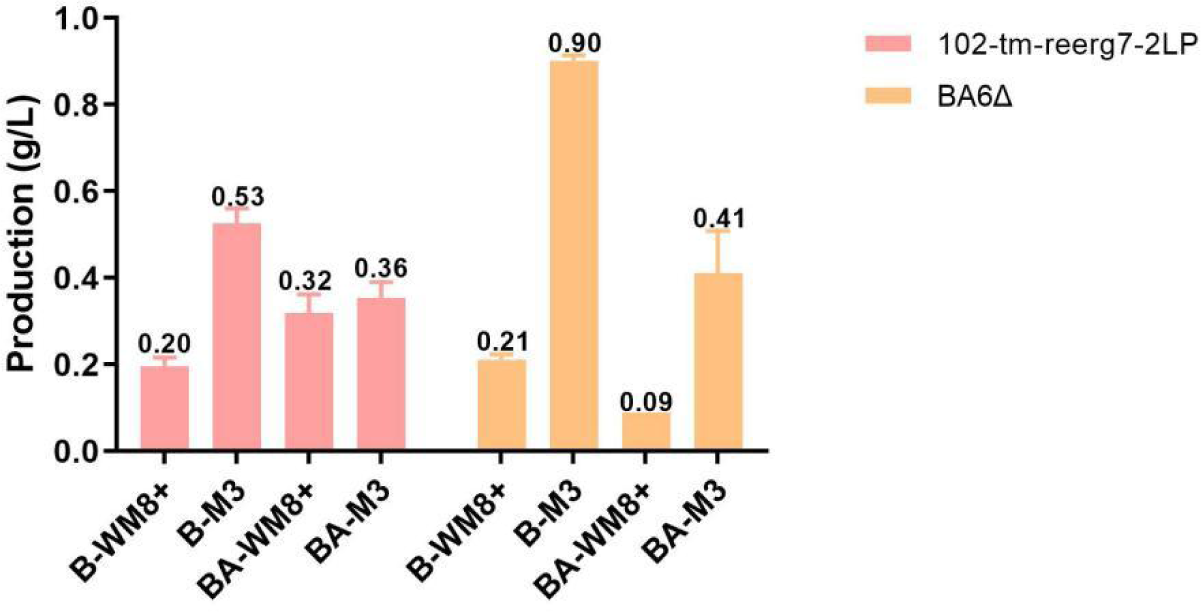
Betulin (B) and betulinic acid (BA) produced by engineered yeasts. The aluminum lid was used for shake flask cover and the media WM8+ and M3 were compared.

In conclusion, we identified the optimum temperature for PPD production at 28 ℃. The M3 medium presented here for producing PPD by yeast was built by stepwise evaluation and optimization of the WM8+ medium. Furthermore, we used a membrane lid to improve the final PPD production, which facilitated increased aeration. Eventually, a remarkable titer of 1.2 g/L PPD was achieved in the 500 mL shake flask, a 7.5-fold improvement compared to the unoptimized standard WM8+ medium with original fermentation conditions (0.16 g/L). As the medium and growth conditions are easily established and transferable to other triterpenoid-producing bakeŕs yeast, the study might facilitate the simple evaluation of engineered strains in microtiter plates and shake flask cultivations.

## MATERIALS AND METHODS

### Strains, Medium, and Reagents

The engineered yeast 102-tm-reerg7-2PP for PPD synthesis was modified from *S. cerevisiae* CEN.PK 102-5B and originated from a previous study.^16,36^ *S. cerevisiae* BA6Δ (unpublished) is a pah1 deletion mutant of BA6, generated through replacement of PAH1 with a hygromycin resistance cassette (YMR165CΔ:hphMX).^37,38^ The engineered yeast BA6Δ for betulin synthesis was modified from *S. cerevisiae* Simo1575.^16^ Standard WM8+ medium included 50 g/L glucose, salt solution, 10 g/L glutamate, trace elements solution, vitamins solution, verduyn-trace elements solution, and verduyun-vitamins solution. In addition, the salt solution includes 0.25 g/L (NH_4_)H_2_PO_4_, 2.8 g/L NH_4_Cl, 0.25 g/L MgCl_2_·6H_2_O, 0.1 g/L CaCl_2_·2H_2_O, 2 g/L KH_2_PO_4_, 0.55 g/L MgSO_4_·7H_2_O, and 0.075 g/L C_6_H_12_O_6_. The trace elements includes 1.75 mg/L ZnSO_4_·7H_2_O, 0.5 mg/L FeSO_4_·7H_2_O, 0.1 mg/L CuSO_4_·5H_2_O, 0.1 mg/L MnCl_2_·4H_2_O, 0.1 mg/L Na_2_MoO_4_·2H_2_O, and EDTA (0.04 mM). The vitamins solution includes 10 mg/L C_6_H_5_NO_2_ (nicotinic acid), 25 mg/L C_8_H_11_NO_3_ (pyridoxine), 10 mg/L C_12_H_17_N_4_OS (thiamine), 2.5 mg/L C_10_H_16_N_2_O_3_S (biotin), and 50 mg/L C_9_H_17_NO_5_ (calcium-d-pantothenate). Verduyn-Trace elements solution includes 15 mg/L Na_2_EDTA, 4.5 mg/L CaCl_2_·2H_2_O, 4.5 mg/L ZnSO_4_·7H_2_O, 3 mg/L FeSO_4_·7H_2_O, 0.3 mg/L CuSO_4_·5H_2_O, 1 mg/L MnCl_2_·2H_2_O, 0.4 mg/L Na_2_MoO_4_·2H_2_O, 0.3 mg/L CoCl_2_·6H_2_O, 1 mg/L H_3_BO_3_, and 0.1 mg/L KI. Verduyn-vitamins solution includes 25 mg/L C_6_H_12_O_6_ (myo-inositol), 1 mg/L C_6_H_5_NO_2_ (nicotinic acid), 1 mg/L C_8_H_11_NO_3_ (pyridoxine), 1 mg/L C_12_H_18_Cl_2_N_4_OS (thiamin-HCl), 0.05 mg/L C_10_H_16_N_2_O_3_S (biotin), 1 mg/L C_9_H_17_NO_5_ (calcium-pantothenate), and 0.2 mg/L C_7_H_7_NO_2_ (p-aminobenzoic acid).^10^ The PPD standard was purchased from PhytoLab. The DM standard was purchased from AOBIOUS.

### Design of Experiment

The 34 runs DoE was designed using Design Expert 11 (Stat-Ease, Inc.). Study type: Factorial; Subtype: Randomized; Design type: Minimum Run Resolution V; 34 runs; Design Model: 2FI; Blocks: No Blocks; Center Points: 4. The experiments were carried out in 24-well plates (polypropylene square deep well MTP, re-usable, 24 wells of 11 ml volume, flattened, detoxified, Kuhner shaker, Germany).

### Ordinary least squared (OLS) Estimation

An OLS regression was used with two sets of linear formulas: 1. containing the individual factor (∑ c_i_F_i_, with c_i_ as the coefficient for each factor F_i_=[A, B, C, D, E, F, G]) and 2. the individual factor with first order factor interactions (∑c_i_F_i_ + ∑c_ij_F_i_F_j_, with c_ij_ as the coefficient for each two-factor interaction). A regression was performed with the seven factors A-G towards either PPD or DM, resulting in four regressions. All data was centered to have a mean value of 0 and a standard deviation of 1. The evaluation of the OLS models is based on the F-statistics, which reports whether a combination of independent variables (the factors in the OLS model) can explain the variance of the dependent variable (PPD or DM) if the p-value is below 0.05. The factors are selected by their individual p-values of their contribution to the OLS model. The impact of each factor is then determined by the respective coefficient. The calculations are performed with the *ols* package in the *statsmodel* (v0.13.2) library in *Python*, and the code is provided as Jupyter Notebook in the Supplementaries.

### Cultivation of Engineered Yeasts in Shake Flasks

Pre-cultures in the 500 mL shake flasks with an aluminum lid containing 50 mL WM8+ medium and inoculate each pre-culture with 500 µL of a thawed cryogenic stock of the engineered yeast (30 ℃, 200 rpm, and 50 mm shaking diameter). The pre-culture medium with an OD_600_ of 6-10 was used as the inoculum of the main culture. The start OD_600_ of the main culture was set to 0.4, measured with a single-use cuvette (polystyrene, semi-micro, 1.6 mL, Carl Roth, Germany) in a photometer (Biochrom Ltd., United Kingdom). All experiments were carried out in the 500 mL shake flasks with aluminum or membrane lids. The filling volume differed between 25 and 50 mL (see text for details). The cultivation temperature differed between 28 °C and 30 °C (200 rpm and 50 mm shaking diameter). At least duplicates were performed.

### Determination of PPD, DM, and Biomass concentrations

800 µL of cell broth were sampled, transferred into 2 mL reaction tubes (Eppendorf, Germany), and frozen at −20 ℃ until further use. For cell disruption, 250 µL of glass beads (0.5 mm diameter, Carl Roth, Germany) were added, followed by 800 µL chloroform: methanol (80:20 ratio) and 80 µL HCl (1 M). The samples were then placed into the MinibeadBeater (Biospec Products, USA) for 2 minutes twice. For phase separation, samples were centrifuged for 10 minutes at 4 ℃, 13,000 rpm (Eppendorf Centrifuge 5418 R, Eppendorf, Germany). 500 µL of the lower organic phase containing the PPD and DM were filtered (4 mm syringe filters RC membrane, pore size 0.2 µm, Phenomenex, Germany) and transferred into HPLC vials (Flat N9 vials, screw neck, 1.5 mL, clear, Macherey-Nagel, Germany) and stored at 4 ℃ until HPLC analysis. PPD and DM concentration were determined with an HPLC system (UltiMate 3000 HPLC system, Thermo Fisher Scientific, USA) coupled to a Dionex Corona Veo Charged Aerosol Detector. The column was an EC150/4.6 NUCLEODUR C18 gravity column (Macherey-Nagel GmbH and Co. KG, Germany), the mobile phase was Acetonitrile:0.2 % (v/v) formic acid, the column temperature was set to 40 ℃, the flow rate was programmed to 1.2 mL/min, the injection volume was set to 5 µL, the runtime was 23 min. To measure the CDW, 2 mL culture was harvested by centrifugation for 5 minutes at 4 ℃, 13,000 rpm (Eppendorf Centrifuge 5418 R, Eppendorf, Germany). The collected yeast cells were washed with deionized water and oven-dried to a constant weight. An OD_600_ to CDW correlation of CDW (g/L)=0.2×OD_600_+1.3 was estimated. All values are reported as the average of three replicates.

### Water evaporation

Water evaporation was monitored using 500 mL shake flasks with either the aluminum or membrane lid. Different water volumes (25 mL and 50 mL) were incubated in a shaker for 120 h (28 ℃, 200 rpm, and 50 mm shaking diameter).

## Author Contributions

Shangkun Qiu completed all the experiments, including the DoE design, OLS estimation analysis, triterpenoids measurement, and data analysis. Shangkun Qiu also wrote the first draft of the manuscript. Mariam Dianat Sabet Gilani helped with the yeast fermentation and triterpenoid measurements. Conrad Müller advised on the DoE experiment design and DoE data analysis. Ulf Liebal advised the OLS estimation analysis. Lars M. Blank supervised the study, discussed the results, and revised the manuscript.

## Declaration of Competing Interest

The authors declare that they have no known competing financial interests or personal relationships that could have appeared to influence the work reported in this paper.

## Acknowledgments

Shangkun Qiu was funded by the China Scholarship Council (CSC). The China Scholarship Council is the Chinese Ministry of Education’s non-profit organization that provides support for international academic exchange with China. Lars M. Blank acknowledges funding from the European Union’s Horizon 2020 Research and Innovation Program under Grant Agreement 870294 for the EU/China Project MIX-UP.

## Notes

### Competing Interest Statement

The authors have declared no competing interest.

### Summary of Updates

Keywords updated.

