## Supplementary Material for "Bioprocess optimization enables enhanced protopanaxadiol production in yeast"

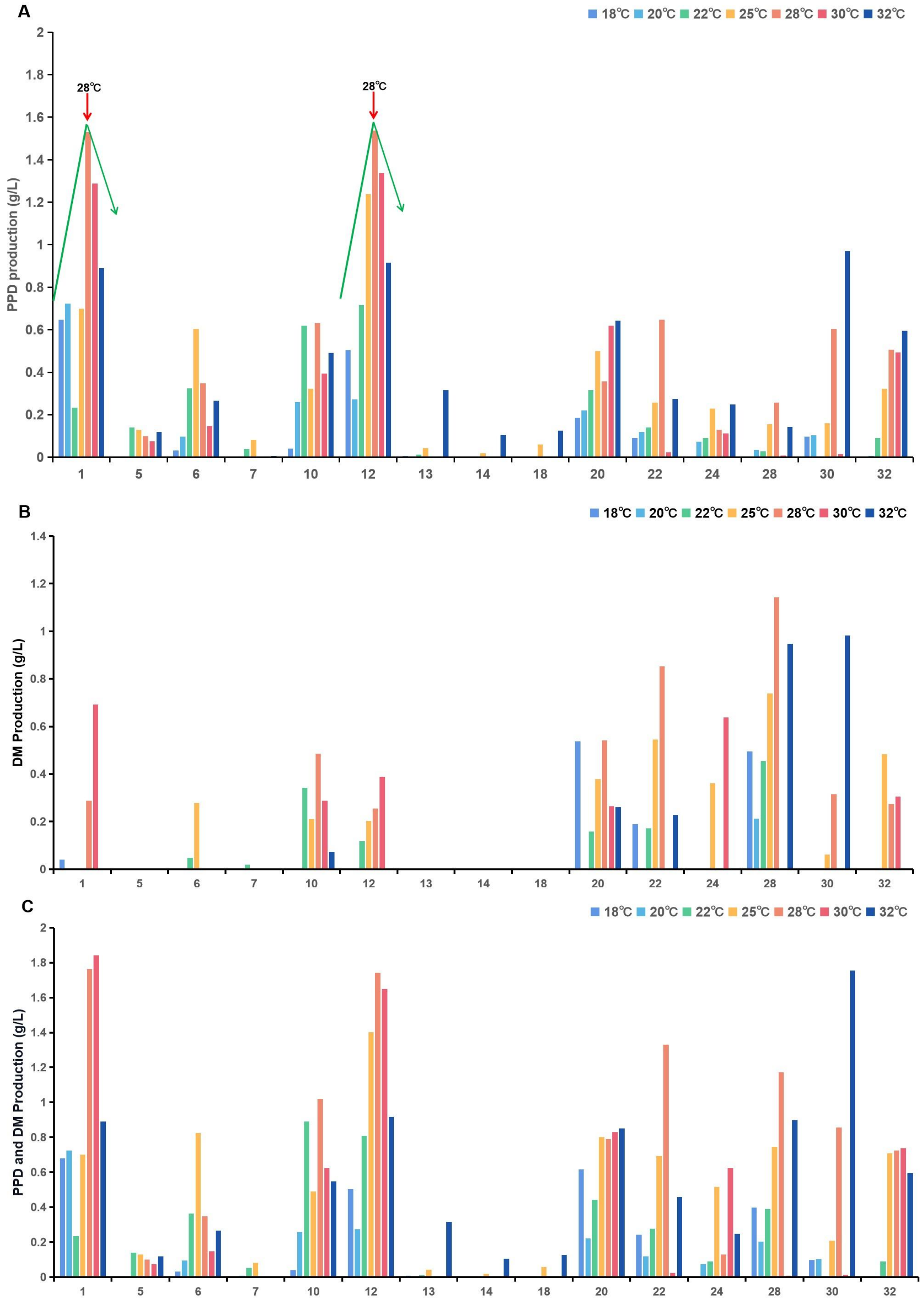


**Figure S1.** The effect of different temperatures on DM and PPD production. (A) The comparison of PPD production at 18℃, 20℃, 22℃, 25℃, 28℃, 30℃, and 32℃. (B) The comparison of DM production at 18℃, 20℃, 22℃, 25℃, 28℃, 30℃, and 32℃. (C) The comparison of total triterpenoid production at 18℃, 20℃, 22℃, 25℃, 28℃, 30℃, and 32℃. The horizontal coordinate is every run's number of the DoE experiment. The medium composition of these runs are same with Table 1.
